# A high-throughput strategy for enhancing aptamer performance across different environmental conditions

**DOI:** 10.1101/2022.09.13.507853

**Authors:** Leighton Wan, Alex Yoshikawa, Michael Eisenstein, H. Tom Soh

## Abstract

Aptamers selected under specific environmental conditions (*e*.*g*., pH, ion concentration, temperature) often exhibit greatly reduced affinity when used in other contexts. This can be especially problematic for biomedical applications, in which aptamers are exposed to sample matrices with distinctive chemical properties, such as blood, sweat, or urine. We present a high-throughput screening procedure for adapting existing aptamers for use in samples whose chemical composition differs considerably from the original selection conditions. Building on prior work from our group, we have utilized a modified DNA sequencer to screen more than 10^6^ aptamer single and double mutants for target binding under the desired assay conditions. As an exemplar, we screened mutants of a previously reported glucose aptamer that was originally selected in high-ionic strength buffer and exhibits relatively low affinity under physiological conditions. After a single round of screening, we identified aptamer mutants with ∼4-fold increased affinity under physiological conditions. Interestingly, we found that the impact of single-base substitutions was relatively modest, but observed considerably greater binding improvements among the double mutants, highlighting the importance of cooperative effects between mutations. This approach should be generalizable to other aptamers and environmental conditions for a range of applications.

## Introduction

Aptamers are nucleic acid-based affinity reagents that are generated based on their target-binding properties through *in vitro* screening using systematic evolution of ligands by exponential enrichment (SELEX).^1,2^ These selection protocols are typically designed to generate an enriched aptamer pool that exhibits high affinity against a target in a defined set of environmental conditions such as pH, ion concentration, and temperature. As a consequence, aptamers often exhibit greatly reduced binding performance when used in conditions that are notably different than those used during selection^3,4^. For example, the concentration of divalent cations such as Mg^2+^ is crucial for the formation of aptamer tertiary structure, including binding pockets responsible for target recognition^5–8^. In another example, an aptamer that binds tetracycline with high affinity in the presence of Mg^2+^ loses its ability to bind this target in the absence of magnesium^5^. pH and Na^+^ ion concentrations can likewise influence ligand binding through mechanisms such as protonation of aptamer binding sites. A cocaine-binding aptamer that exhibited an optimal K_D_ at pH 7.4 lost binding function at pH 9.6, along with a 20-fold reduction in affinity at increased NaCl concentrations^9^. This becomes a critically important consideration in the context of real-world applications such as molecular diagnostics. A high-quality aptamer selected in a particular buffer may lose affinity or specificity when used in clinically relevant specimens with chemical properties that differ meaningfully from that buffer, such as blood, sweat, or urine, requiring the identification of new aptamers that perform well in the desired conditions.

We describe here a high-throughput mutational screening procedure for adapting existing aptamers for use in assay conditions that differ from those used in the initial selection. This strategy employs our previously published non-natural aptamer array (N2A2) platform,^10,11^ making use of a modified Illumina MiSeq instrument to screen the binding properties of vast pools of single and double substitution mutants of the parent aptamer sequence. As a proof of concept, we applied this approach to a previously published glucose aptamer, which exhibits an equilibrium dissociation constant (*KD*) of 10 mM when used in the high-salt HEPES-based buffer employed in the original SELEX process.^12^ We found that this aptamer exhibits a nearly 10-fold higher *K*_*D*_ (*i*.*e*., lower affinity) at physiological salt concentrations, and this would make it less useful for detecting physiological concentrations of blood glucose, which typically range between 5.6 mM during fasting and 7.8 mM after eating in healthy individuals.^13^ Using our platform, we were able to discover mutant versions of this aptamer that exhibit nearly four-fold improved affinity for glucose in these lower-salt conditions, offering a better match for physiological applications. Since our platform enables comprehensive screening of single- and double-mutant libraries for a given aptamer, we were also able to identify key structural features of the aptamer that could offer fruitful targets for further engineering and functional optimization of the aptamer sequence.

## Materials and Methods

### Reagents

The mutant library, which contained single and double substitutions at every position of the base glucose aptamer and was flanked by Illumina adapter sequences with an additional EcoRI cutsite, was synthesized by TWIST Bioscience. All other sequences, including fluorophore (Cy3)- and quencher (DABCYL)-modified candidate sequences and displacement strands, were ordered HPLC-purified from Integrated DNA Technologies (IDT), along with IDTE (pH 8.0) buffer. All sequences used in this work are listed in **Supplementary Table 1**. Sequencing reagents including the Nextera Index Kit (#15055290) and 150-cycle MiSeq Reagent Kit v3 (#MS-102-3001) were ordered from Illumina. Library sequencing preparation was performed using the GoTaq Master Mix from Promega (#M7433). D-(+)-Glucose was ordered from Sigma-Aldrich (#G8270). Corning 96-well half-area black flat-bottom polystyrene microplates (Thermo Fisher Scientific, No. 07-201-205) were used in the plate reader assays. All buffers and chemicals were ordered from Thermo Fisher Scientific unless specified otherwise.

### High-throughput sequencing and screening preparation

The mutant library was resuspended in IDTE buffer to 20 ng/µL. The Nextera Index Kit and the GoTaq Master Mix was used to index 90 ng of the pool in a 450 µL PCR reaction volume, following the manufacturers’ instructions. We performed 10 cycles of amplification based on a pilot PCR protocol using 50 µL of the overall sample, where 4 µL was collected every other cycle. PCR cycles consisted of 30 seconds at 95 °C, 30 seconds at 55 °C, and 30 seconds at 72 °C. The Axygen AxyPrep Mag PCR Clean-Up Kit (#MAG-PCR-CL-50) was used to clean the final product before verification on a 10% TBE gel.

### High-throughput mutational screen

A modified Illumina Miseq and high-throughput screening platform developed by our group was used to sequence and screen ∼10^7^ aptamer clusters, as described in previous work^10,11^. Briefly, monoclonal aptamer clusters containing ∼1,000 strands per cluster were generated during canonical bridge amplification. Afterwards, during the first read, sequences and locations on the flow-cell were determined. Before measuring aptamer cluster binding to the displacement strand and targets, the reverse primer adapter sequence was removed using EcoRI (New England Biolabs, #R0101L) and strand complementary to EcoRI and part of the adapter sequence (**Supplementary Table 1**). During the buffer cycle, the flow-cell was first cleaned with 750 µL of 0.05 M NaOH with 0.25% SDS and then washed with 500 µL of buffer. We then introduced 515 µL of 0.2 µM Cy3-labeled displacement strand and heated to 80 °C before ramping down to 22 °C over ∼30 minutes. Excess strands were removed by washing with 6 mL of buffer. The flow-cell was then imaged to conclude the buffer cycle. During the target cycle, the flow-cell was washed with 1.25 mL of buffer before adding 100 mM glucose in buffer. The HEPES buffer consisted of 20 mM HEPES, 1 M NaCl, 10 mM MgCl_2_ and 5 mM KCl, with an ionic strength of 1.035 M. Selection buffer consisted of 20 mM Tris-HCl, 120 mM NaCl, 5 mM KCl, 1 mM MgCl_2_, 1 mM CaCl_2_, and 0.01% Tween 20, with an ionic strength of 0.144 M. 500 µL of the 100 mM glucose solution was then added, followed by seven more additions of 15 µL of 100 mM glucose solution, with five-minute incubations between each addition. Measurements were made in triplicate for both glucose and no-glucose control cycles. Glucose-free ‘no-target’ cycles were used as a control to account for displacement during target cycles due to washing with buffer. Sequence-intensity data was linked from Miseq FASTQ, cif, and loc files using custom Python code (https://github.com/sohlab/non-natural-aptamer-array).^10^

### Data processing and mutant mapping

The sequence-intensity data were first filtered before mapping and identification of aptamer candidates. Clusters with intensities < 80 RFU in buffer cycles were removed to account for system noise, and clusters with intensities >3,000 RFU in any cycle were removed to account for quantification errors or imaging artifacts. To normalize for noise resulting from experimental or imaging differences in each cluster, the intensities for pairs of buffer and subsequent target cycles were converted to percent changes. A greater percent change corresponded to a larger drop in intensity during the target cycle. The percent change for glucose cycles was normalized by subtracting the percent change in no-target cycles. Afterwards, the data were filtered to remove sequences represented by <5 clusters after initial filtering for consistency. Sequences were ranked by mean normalized percent change to find the top performing sequences for subsequent analysis.

### Plate reader characterization

We used a previously-published plate-reader assay^12^ to measure aptamer affinity for the displacement strand (K_D,1_) and the affinity of the target for the aptamer-displacement strand complex (K_D,2_). The K_D_ is equal to the ratio of K_D,1_/K_D,2_. K_D,1_ was measured by titrating a DABCYL-labeled displacement strand with 50 nM Cy3-labeled aptamer in buffer. Strands of various lengths were tested to find a displacement strand that achieved 90% quenching using 100–500 nM quencher strand. The 13-mer strand achieved this goal for each aptamer candidate and was used for subsequent testing. The assay was performed by incubating 50 nM Cy3-labeled aptamer with the displacement strand in buffer at 95 °C for 5 minutes, before cooling down to room temperature for at least 30 minutes. The final reaction volume (100 µL) was measured at room temperature (25 °C) in opaque black half-well plates on a Synergy H1 microplate reader (BioTek) with a filter cube (emission: 590/35, excitation 540/63) at gain of 55. The fluorescence data for the displacement strand assays were fit using Python, with the ‘curve_fit’ function from the ‘scipy’ library. Data were fitted using the Hill equation, 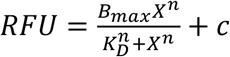 with *n* = 1. The concentration of displacement strand was then determined for the subsequent assay with target for each strand. We used 250 nM displacement strand for glu0, glu2, G24A, and T30C, and 125 nM for glu1 and T30C/A41T. K_D,2_ was then determined using 50 nM Cy3-labeled aptamer, the previously identified concentration of 13-mer displacement strand, and 0 to 1600 mM glucose in buffer. Before fitting, the data was normalized to account for the glucose concentration-dependent increase in fluorescence observed at higher concentrations (**Supplementary Figure 4A**). To normalize the increased fluorescence, we first measured the fluorescence of 50 nM Cy3-labeled 14-mer control strand (**Supplementary Table 1**) at the same concentrations of glucose (0 to 1600 mM) in buffer. The ratio of fluorescence increase at a given concentration was then calculated by dividing the control strand fluorescence measured at the chosen concentration of glucose by the control strand fluorescence measured without glucose. The fluorescence measured with aptamer-displacement strand complex plus glucose was then divided by the ratio of fluorescence increase from the control strand at the corresponding concentration. Each measurement was made in triplicate. These data were then fitted and used to calculate K_D,2_, after which we determined the K_D_.

## Results

### Design of the high-throughput mutant screening process

We used our N2A2 platform as a means for comprehensively surveying the structure and function of the full landscape of single and double mutants for our glucose aptamer (glu0), which was initially isolated by Nakatsuka *et al*.^12^ N2A2 is based on a modified Illumina Miseq sequencing instrument, which performs both high-throughput sequencing and screening of aptamer clusters on the flow-cell surface. The MiSeq flow-cell can accommodate as many as 10^7^ aptamer clusters, which is more than sufficient to achieve full coverage of the single and double mutant space for most existing aptamers in a single experiment—for example, the sequence space of single and double substitution mutations for an 80-mer aptamer consists of 28,681 sequences.

The canonical first read of the high-throughput sequencing run is used to generate and sequence the aptamer clusters (**Figure 1A**). Instead of a traditional second, paired-end read, the N2A2 then measures the fluorescence of the aptamer clusters, which is produced through the hybridization of a fluorescently-labeled, complementary displacement strand. This strand is displaced when the aptamers at a given cluster bind to the target molecule, providing a measurable readout of binding activity. This activity is measured in alternating ‘buffer’ and ‘target’ cycles (**Figure 1B**). During the buffer cycle, the fluorescently-labeled displacement strand is annealed to aptamer clusters, washed with buffer, and then imaged to measure the affinity for the displacement strand. In the following target cycle, the flow-cell is incubated with the target (*i*.*e*. glucose) in buffer, and then washed and imaged in order to measure the extent to which displacement strand signal is reduced relative to the buffer cycle (**Figure 1C**). A large relative difference indicates a greater response to the target, presumably due to binding-induced strand displacement, whereas a smaller or no fluorescence change generally indicates aptamers that have minimal target binding activity. In this fashion, the relative degree of aptamer binding to targets can be measured while linking this binding performance directly to the relevant aptamer cluster sequence. Finally, the response to target can be used to generate a mutation map to investigate aptamer function and sequence elements that can potentially be targeted to achieve improved binding performance (**Figure 1D**).

**Figure 1:**
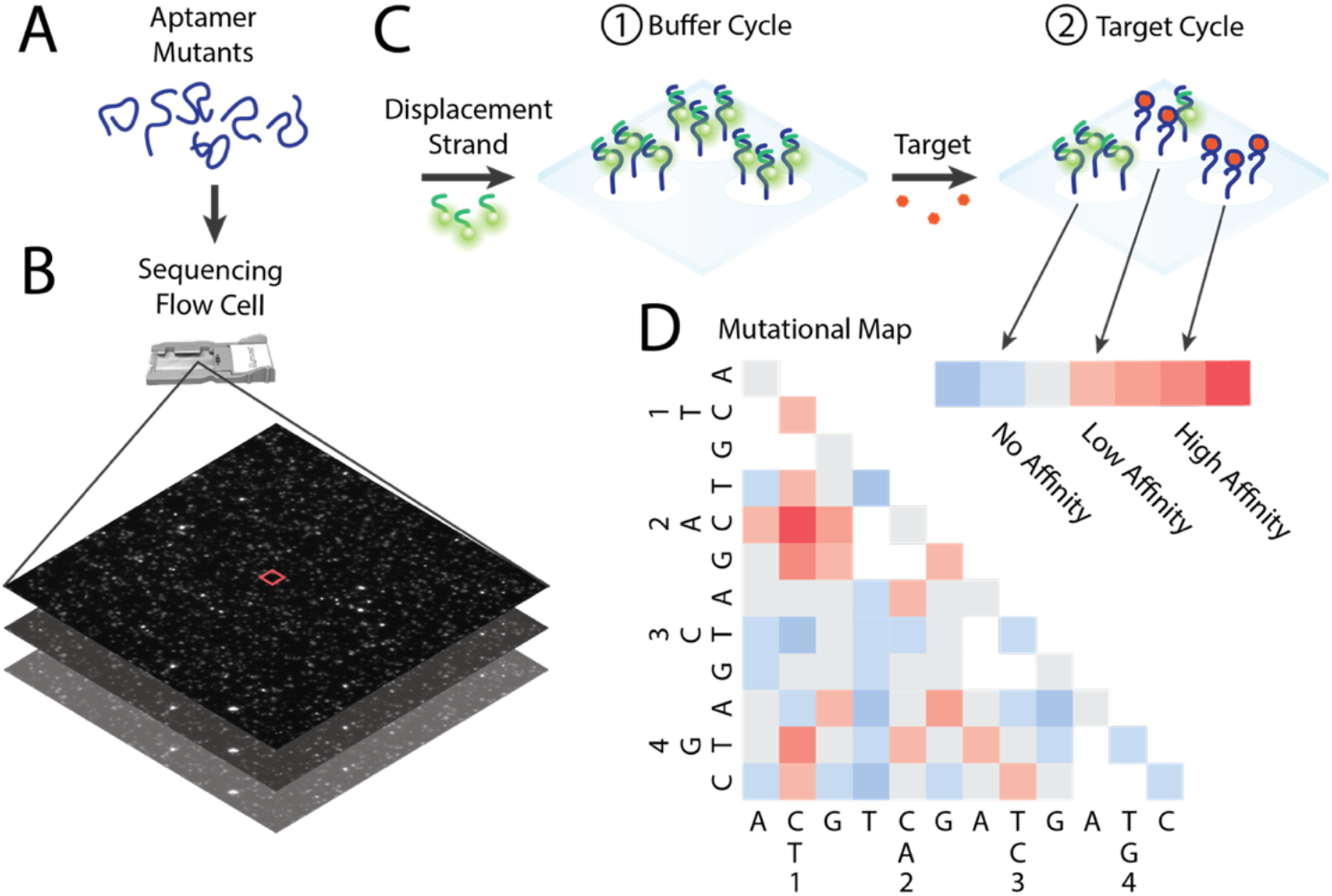
Workflow for high-throughput screen of aptamer mutants for binding optimization in new buffer. **A)** Aptamer mutants are converted to aptamer clusters and then screened using a modified Illumina sequencer and **B)** flow-cell. **C)** The screen is performed using alternating ‘buffer’ and ‘target’ cycles. During buffer cycles, Cy3-labeled displacement strand is added, washed in selection buffer, and then the clusters are imaged. During target cycles, 100 mM target in the same buffer is added, followed by washing and imaging. The process is repeated for both the original and desired selection buffer. **D)** The high-throughput screening data is then used to generate a mutational map from which candidate sequences are chosen for further characterization.

### Mutational screen of glucose aptamer response in physiological conditions

The glu0 aptamer tested here was originally selected in a high-ionic-strength (1.035 M) HEPES buffer, and exhibits a *K*_*D*_ of 10 mM in this buffer. However, we found that its affinity is greatly reduced in our selection buffer (20 mM Tris-HCl, 120 mM NaCl, 5 mM KCl, 1 mM MgCl_2_, 1 mM CaCl_2_, and 0.01% Tween 20), which more closely mirrors physiological conditions with an ionic strength of 144 mM. We measured the aptamer’s *K*_*D*_ in both HEPES buffer and selection buffer using the same plate-reader assay used in the original paper. Briefly, this assay measures the *K*_*D*_ of the aptamer as the ratio of the aptamer’s affinity for a displacement strand (*K*_*D,1*_) relative to the response of the aptamer-displacement strand complex to the target (*K*_*D,2*_) (**Supplementary Figure 1A**). The displacement strand sequence was based on that from the original paper (**Supplementary Table 1**). Based on this assay, we measured a K_D_ of 3.65 mM in HEPES buffer versus 90.5 mM in selection buffer (**Supplementary Figure 1B-C**). As such, this aptamer would be of limited use for measuring physiological levels of glucose in blood, which range from <5.6 mM during fasting and up to 7.8 mM after eating in healthy individuals and are greater in diabetic patients.^13^

In order to profile the binding of various aptamer mutants in our selection buffer, we generated a pool of 11,628 single- and double-mutants based on the 51-mer glu0 sequence. The pool was then indexed and prepared for high-throughput sequencing and screening. During buffer cycles, we measured aptamer cluster binding to a Cy3-labeled 15-mer displacement strand (**Supplementary Table 1**). In the subsequent target cycles, we introduced either 100 mM glucose or target-free buffer as a control, and once again measured fluorescence. We chose a stringent concentration of 100 mM glucose to accommodate for competition with the displacement strand. This is well above the measured K_D_ for glu0 in both selection buffer and HEPES, and should thus readily reveal aptamer variants with superior binding properties. We then processed the sequence-intensity data by filtering out outliers and artifacts and removing sequences with < 5 replicates to minimize the impact from experimental noise (see **Materials and Methods** for details). To control for greater displacement of the Cy3-labeled strand in absence of target, we normalized our measurements by subtracting the percent change in target-free control cycles from the percent change in 100 mM glucose cycles. Hereafter, “percent change” in response to glucose refers to this normalized measurement.

We measured a total of 3.7 million aptamer clusters to identify aptamers with improved affinity for glucose in selection buffer. After filtering, the screened sequences covered 92.0% of the single and double mutant sequence space, with an average of 162 replicate clusters per sequence. We then mapped the mean percent change of each mutant (based on all replicate clusters) onto single- and double-mutant heatmaps to identify mutants that outperformed the parent glucose aptamer (**Figure 2A, B**). Sequences that had minimal or no function would have negative or near-zero percent changes, whereas mutants that retained affinity would have positive percent changes. Overall, 64.5% of the mutants had positive mean percent changes, and 17.4% exhibited a greater mean percent change than that of the base glucose aptamer (2.20%; **Figure 2C**). Among the single mutants, 20.3% of sequences outperformed the base glucose aptamer (glu0), versus 17.3% for the various double mutants, and we subsequently embarked on a closer examination of the effects of specific mutations.

**Figure 2:**
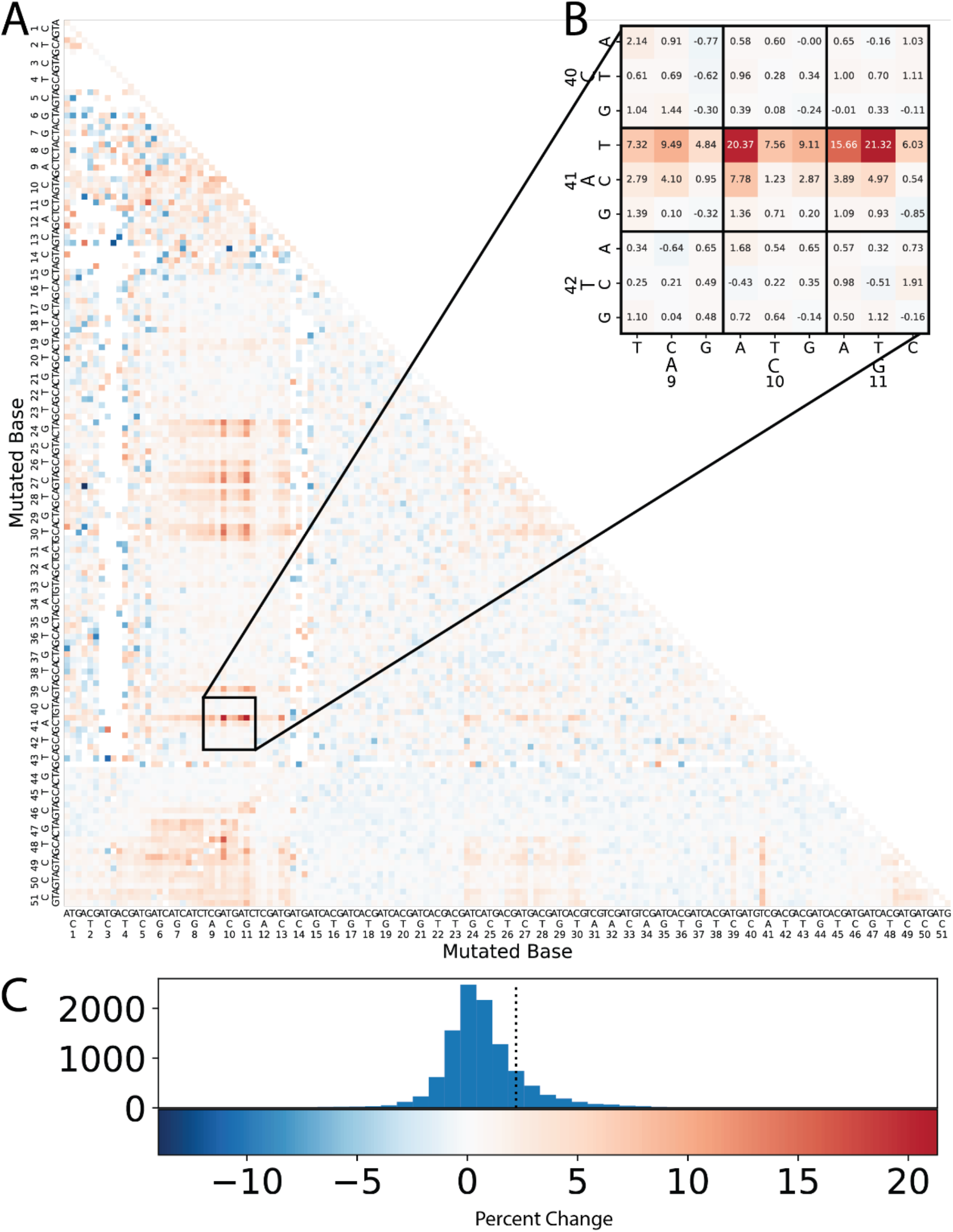
Heatmap of glucose binding by glu0 single and double mutants. **A)** The full sequence map shows all sequences that passed filtering. Colors depict mean normalized percent change values, wherein the percent change observed for a given sequence in selection buffer alone was subtracted from the percent change observed with glucose. Sequences with a large positive percent change are shown in red, while sequences with a negative percent change are shown in blue. Sequences removed in filtering are shown in white. **B)** A magnification of a portion of the heatmap containing the candidate sequences with the highest percent changes. The legend for the heatmap in **A** and **B** is shown at bottom. **C)** The distribution of percent changes for all mutants is shown as a histogram, with the percent change for parent aptamer glu0 marked as a dotted line.

### Characterizing the functional impact of individual aptamer mutations

The high-throughput mutational screen provided insight on specific mutations that influence glucose binding. We categorized the single mutants with mean percent change greater than glu0 into four groups: the displacement strand-binding domain of the aptamer (bases 1-15), the ‘stem-forming’ sequence complementary to part of this domain (bases 44-51), the A41T mutant within the aptamer’s large primary loop, and in a hairpin loop domain (bases 24-30) (**Figure 3A**). Mutations in the displacement strand-binding region of the aptamer are expected to destabilize hybridization with both the displacement strand and the formation of the stem at the end of the aptamer. In contrast, mutations in the complementary stem-forming domain at the end of the aptamer are expected to only destabilize the stem, shifting equilibrium in favor of the aptamer remaining bound to the displacement strand. The single mutants with the highest percent changes were C10A and G11T, with percent changes of 7.59% and 6.82%, respectively. These alterations fell within the displacement strand-binding domain, and other highly-ranked single-substitution sequences also affected this same domain.

**Figure 3:**
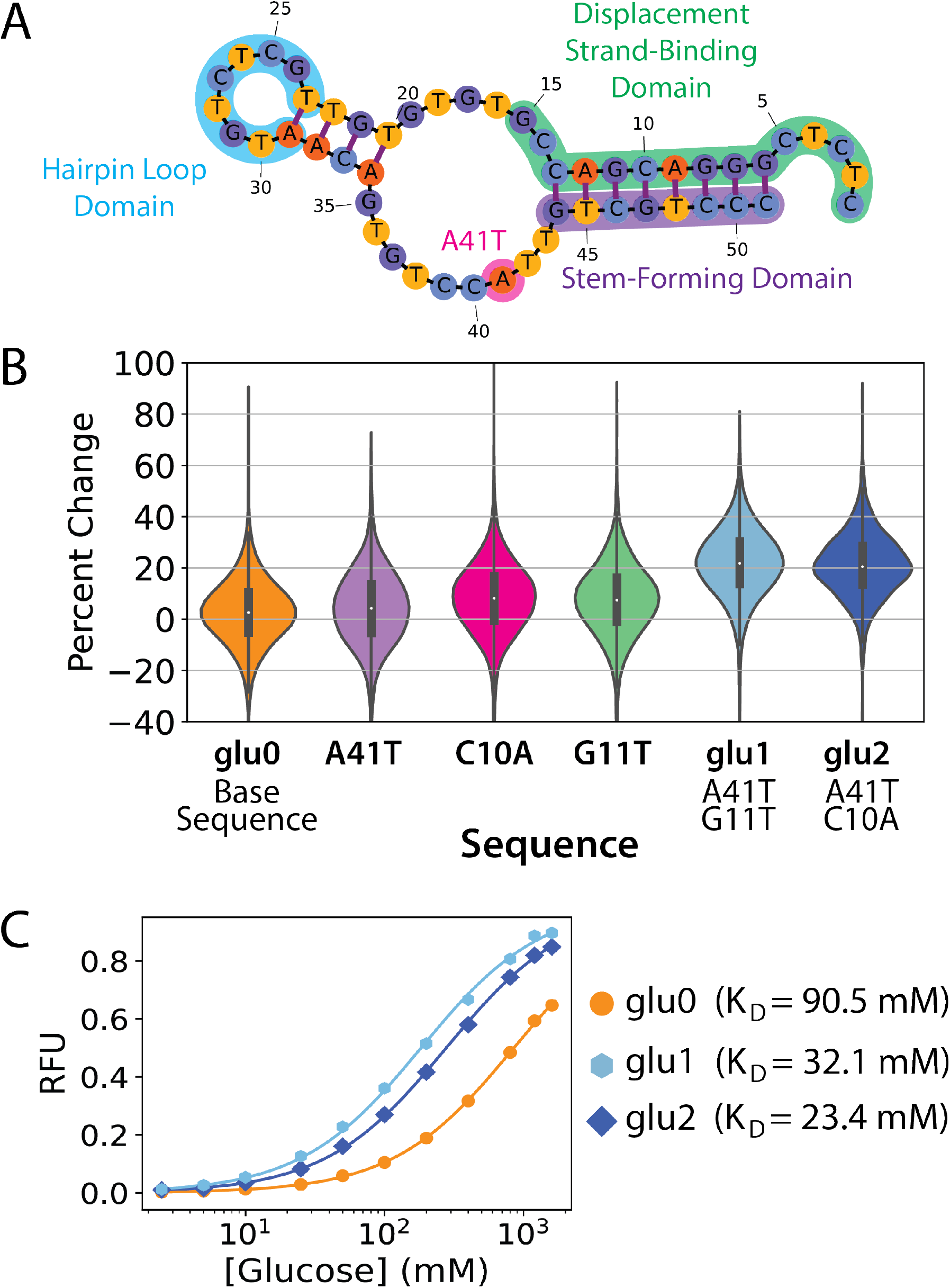
Characterization of high-performing aptamer mutants. **A)** Secondary structure of glu0 as predicted by NUPACK^14^. Four domains are highlighted: the displacement strand-binding domain (green), the hairpin loop (blue), the A41T mutation (magenta), and the stem-forming domain (purple). **B)** Violin plots for the percent changes for the glu0 base sequence and for mutants A41T, C10A, G11T, glu1 (A41T/G11T), and glu2 (A41T/C10A). The thick, dark bar in the center of the violin plot shows the interquartile range, and the white dots mark the median percent change. **C)** K_D_ was measured for glu0, glu1, and glu2 in selection buffer with a plate-reader assay as a ratio of K_D,1_ (**Supplementary Figure 4C**) and K_D,1_ (shown here). The points represent the mean and the bars represent the standard deviation of three measurements.

Although the impact of single substitutions was relatively modest, we observed considerably greater binding improvements in some of our double mutants, highlighting the importance of cooperative effects between mutations. To identify synergistic mutations, we grouped double mutants containing a common single mutation and counted the total number of those mutants with mean percent change greater than glu0 (**Supplementary Figure 2**). The group containing the A41T substitution had the greatest number of double mutants outperforming glu0 (64 sequences), followed by the T30C, C10A, and G24A substitution groups, with 60, 60, and 57 sequences, respectively. Our two highest-performing sequences, A41T/G11T (glu1) and A41T/C10A (glu2), showed percent changes of 21.3% and 20.4%, nearly three times greater than the highest-performing single mutant (**Figure 3B**).

In contrast, while the G24A and T30C groups had high numbers of outperforming mutants, these two mutations did not show synergistic improvements to affinity in combination with other mutants. Individually, the G24A and T30C mutations had low percent changes of 3.00% and 2.91%, respectively, with no notable improvement in affinity (**Supplementary Figure 3A-C**). The T30C/A41T mutant also showed no meaningful cooperativity, with a low percent change of 6.20% and minimal improvement to affinity. In terms of secondary structure, the G24A and T30C mutations are predicted to stabilize the stem preceding the hairpin loop leading to no meaningful impact on glucose binding (**Supplementary Figure 3D**). Since the cooperative effects of mutations cannot currently be determined *a priori*, this high-throughput screening procedure becomes invaluable for systematically investigating how to improve the glucose aptamer through multiple mutations and uncovering structural features that contribute meaningfully to ligand-binding.

### Characterizing mutants with improved glucose response in physiological buffer

We picked the top-performing double-mutant aptamer candidates, glu1 and glu2, for further validation. We used our plate-reader assay to determine the *K*_*D*_ as a ratio of *K*_*D,1*_ and *K*_*D,2*_. Increased fluorescence was measured for a Cy3-labeled 14-mer control strand (**Supplementary Table 1**) at higher concentrations of glucose (**Supplementary Figure 4A**), and we used these background measurements to normalize the measured fluorescence observed from the aptamers with glucose at the same concentrations (**Supplementary Figure 4B**). The calculated K_D_ values for glu1 and glu2 were 32.1 mM and 23.4 mM, respectively, versus 90.5 mM for glu0 in selection buffer (**Figure 3C, Supplementary Figure 4C**), representing a ∼3–4-fold change improvement. This improved K_D_ in selection buffer should yield an aptamer that is more sensitive and applicable to low millimolar concentrations of glucose under physiological conditions. Interestingly, glu1 and glu2 also showed a similar affinity to the parent aptamer when tested in HEPES buffer, with K_D_ values of 3.55 mM and 3.74 mM, respectively (**Supplementary Figure 5**), showing that the mutations identified in the screen broadened the aptamer’s effective operating conditions rather than yielding an aptamer that is solely optimized for use in selection buffer.

## Discussion

In this work, we demonstrate a high-throughput screening strategy that allows us to identify mutant derivatives of an existing glucose aptamer that exhibit improved performance in a physiological buffer relative to the parent sequence, which was originally selected in a high ionic strength buffer. We screened nearly every possible single- and double-mutant of this aptamer for glucose binding in selection buffer using our N2A2 platform. While single substitution mutants achieved a modestly improved response to glucose in selection buffer, we found that certain double mutations yielded synergistic improvements to affinity that allowed them to outperform any of the corresponding single mutants. The two top performing double mutants, glu1 and glu2, exhibited a ∼4-fold improvement over the parent aptamer. Our analysis also yielded a mutation map that could be used to identify areas of the aptamer that might prove useful for further engineering of the aptamer. However, it is also worth noting that the impact of specific mutation combinations on aptamer structure and function was not generally predictable, and this highlights the advantage of using a platform that can broadly screen large numbers of such mutants in a single experiment. We expect that our approach could be applied to other aptamers and environmental conditions to extend the utility of aptamers for a broader set of applications.

