## Supplementary Information for "A high-throughput strategy for enhancing aptamer performance across different environmental conditions"

**Supplementary Table 1:** Sequences used in the high-throughput mutational screen and in the plate reader characterization of sequences.

| Name | Sequence (5' -> 3') |
|---|---|
| Library | *See attached document for mutant pool sequences* |
| Cy3 displacement strand | CGGTCGTCCCGAGAG-Cy3 |
| 13-mer displacement strand DABCYL | CGGTCGTCCCGAG-DABCYL |
| glu0 | Cy3-CTCTCGGGACGACCGTGTGTGTTGCTCTGTAACAGTGTCCATTGTCGTCCC |
| glu1 | Cy3-CTCTCGGGAAGACCGTGTGTGTTGCTCTGTAACAGTGTCCTTTGTCGTCCC |
| glu2 | Cy3-CTCTCGGGACTACCGTGTGTGTTGCTCTGTAACAGTGTCCTTTGTCGTCCC |
| G24A mutant | Cy3-CTCTCGGGACGACCGTGTGTGTTACTCTGTAACAGTGTCCATTGTCGTCCC |
| T30C mutant | Cy3-CTCTCGGGACGACCGTGTGTGTTGCTCTGCAACAGTGTCCATTGTCGTCCC |
| T30C/A41T mutant | Cy3-CTCTCGGGACGACCGTGTGTGTTGCTCTGCAACAGTGTCCTTTGTCGTCCC |
| 14-mer Cy3-labeled control strand | GTCGTCCCGAGAGC-Cy3 |
| EcoRI comp | AGAGACAGGAATTC |

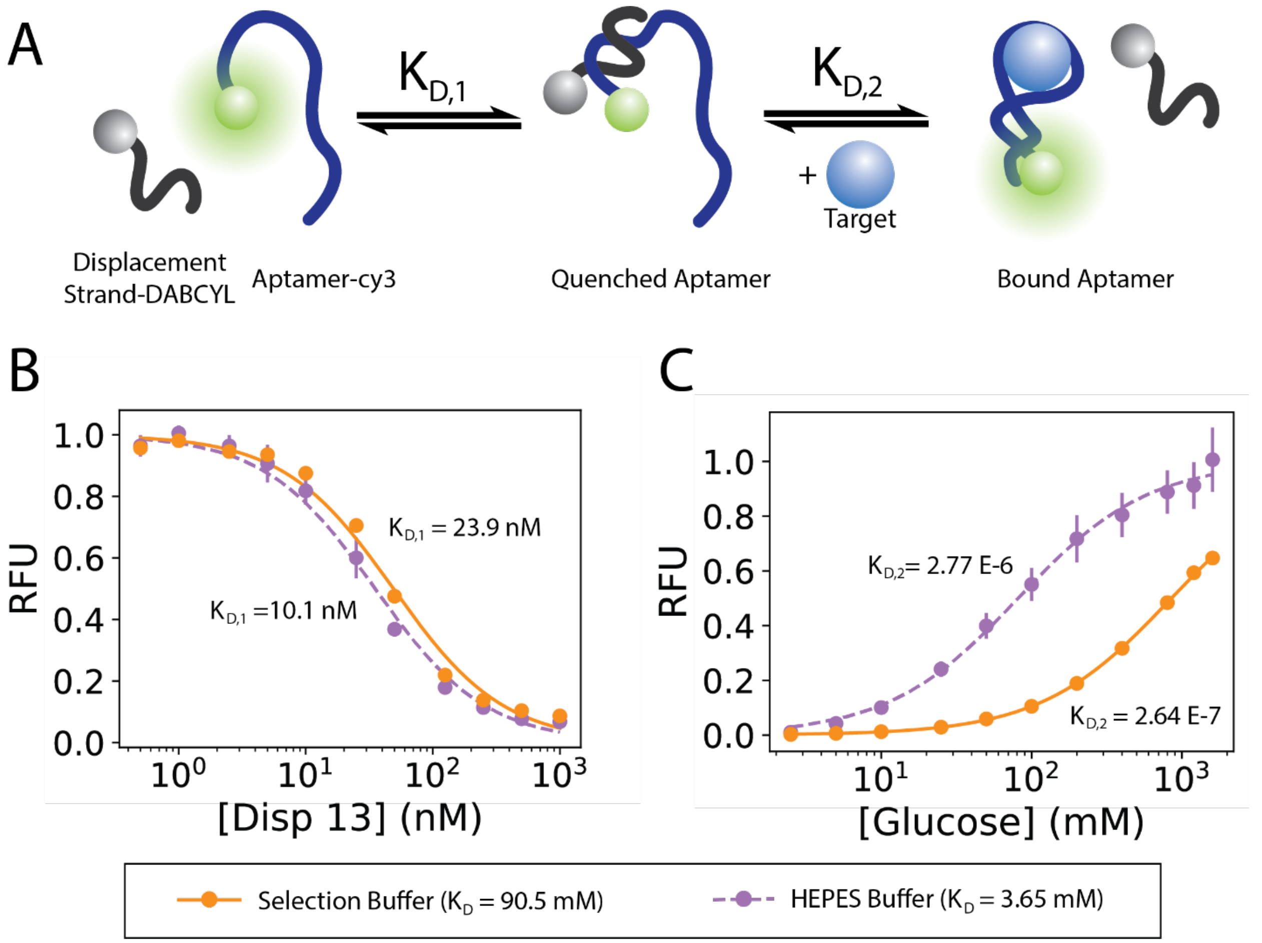


**Supplementary Figure 1**: Measuring aptamer performance using a plate reader assay. **A)** Schematic of the assay. The Cy3-labeled aptamer is first measured for affinity against a DABCYL-labeled displacement strand, as shown in **B**. Afterwards, the quenched aptamer is measured for affinity with the target, as shown in **C**. The final measured $K_D$ values were 90.5 mM and 3.65 mM in selection buffer and HEPES buffer, respectively. The points represent the mean and the bars represent the standard deviation of three measurements.

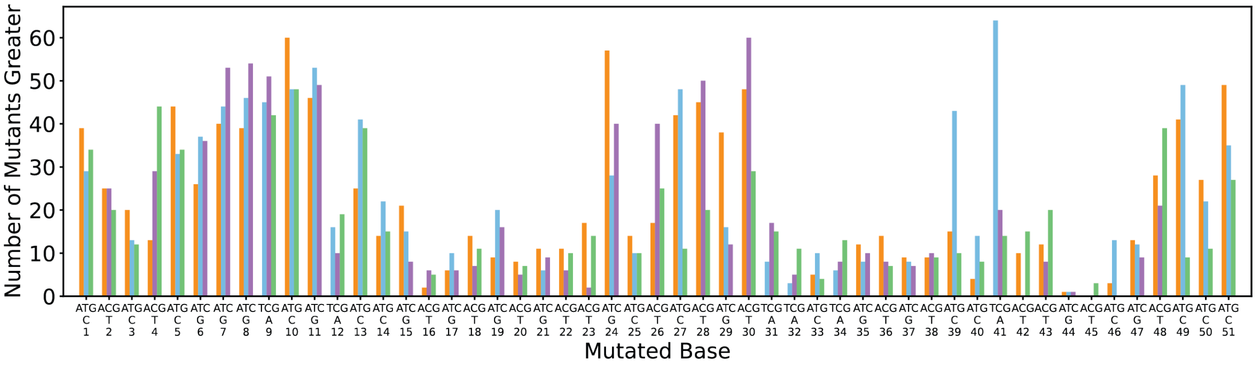


**Supplementary Figure 2**: Identifying cooperative substitution mutations in the double mutant pool. Double mutants sharing a common single mutation were grouped, and we have plotted the number of sequences in each group with a mean percent change greater than that of glu0 (2.20%).

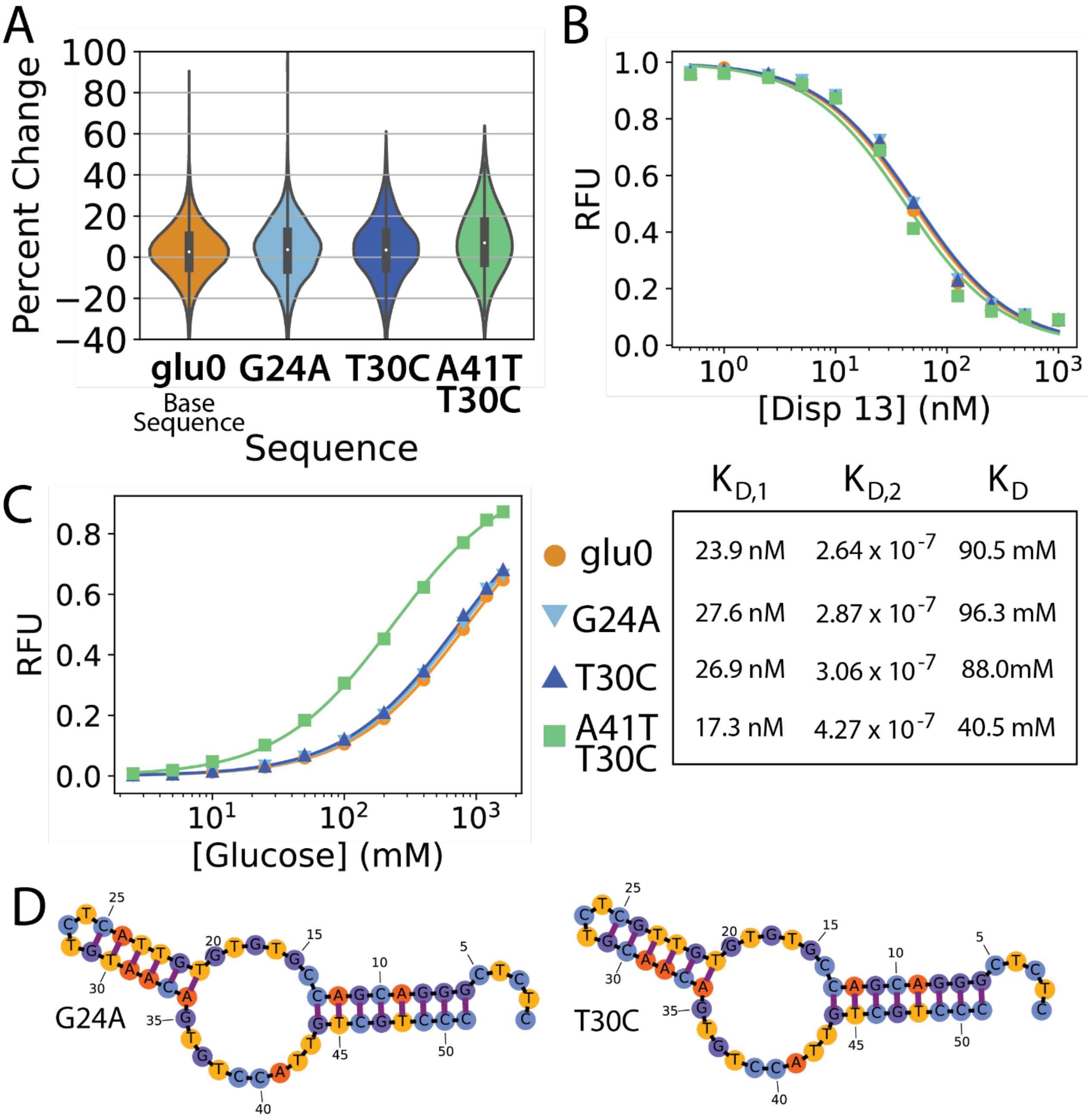


| | $K_{D,1}$ | $K_{D,2}$ | $K_D$ |
|---|---|---|---|
| glu0 | 23.9 nM | $2.64 \times 10^{-7}$ | 90.5 mM |
| G24A | 27.6 nM | $2.87 \times 10^{-7}$ | 96.3 mM |
| T30C | 26.9 nM | $3.06 \times 10^{-7}$ | 88.0mM |
| A41T T30C | 17.3 nM | $4.27 \times 10^{-7}$ | 40.5 mM |

**Supplementary Figure 3**: Non-cooperative single mutations. **A)** Percent changes from glu0 and the G24A, T30C, and T30C/A41T mutants, shown as violin plots. The thick, dark bars in the center of the violin plot show the interquartile range, and the white dots mark the median percent change. **B, C)** We then calculated $K_D$ for G24A (light blue inverted triangle), T30C (dark blue triangle), and T30C/A41T (green square). The points represent the mean and the bars represent the standard deviation of three measurements. **B)** Binding to the 13-mer displacement strand was used to calculate $K_{D,1}$ values. **C)** Aptamer bound to the displacement strand was titrated with glucose to calculate the $K_{D,2}$ values. Final calculations are shown in the table at right. **D)** Secondary structures predicted by NUPACK are shown for the G24A and T30C mutants.

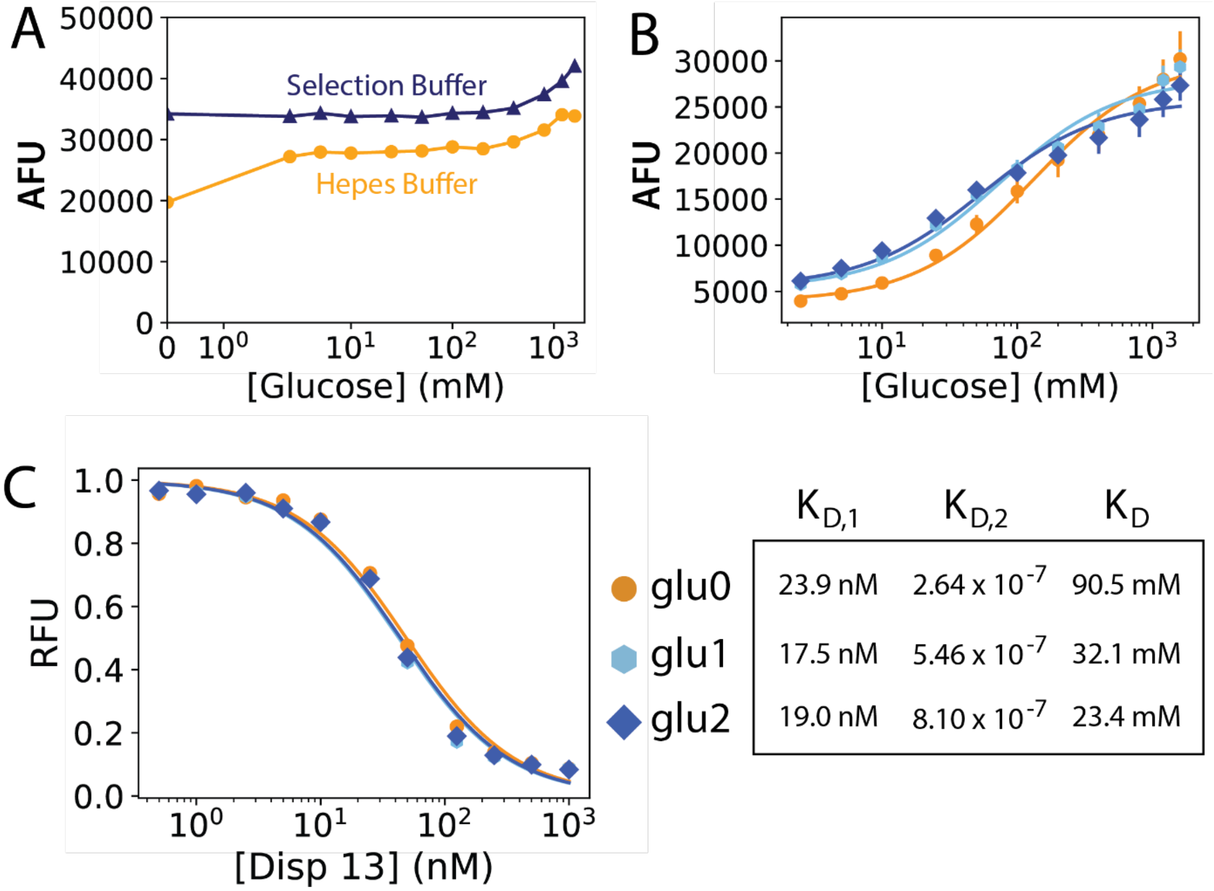


| | $K_{D,1}$ | $K_{D,2}$ | $K_D$ |
|---|---|---|---|
| glu0 | 23.9 nM | $2.64 \times 10^{-7}$ | 90.5 mM |
| glu1 | 17.5 nM | $5.46 \times 10^{-7}$ | 32.1 mM |
| glu2 | 19.0 nM | $8.10 \times 10^{-7}$ | 23.4 mM |

**Supplementary Figure 4**: Measuring the $K_D$ for glu1 and glu2. **A)** Fluorescence measurements from 50 nM Cy3-labeled 14-mer control strand at various concentrations of glucose were used to control for increase in background Cy3 fluorescence at higher concentrations of glucose. We used the ratio of fluorescence at each concentration relative to the fluorescence with no glucose to normalize the fluorescence measurements of each aptamer with glucose. **B)** The unnormalized fluorescence values for glu0, glu1, and glu2. **C)** $K_{D,1}$ for glu0, glu1, and glu2 was assessed based on measurements of aptamer binding to a 13-mer displacement strand. The table at right shows the final calculated $K_D$ values. The points represent the mean and the bars represent the standard deviation of three measurements.

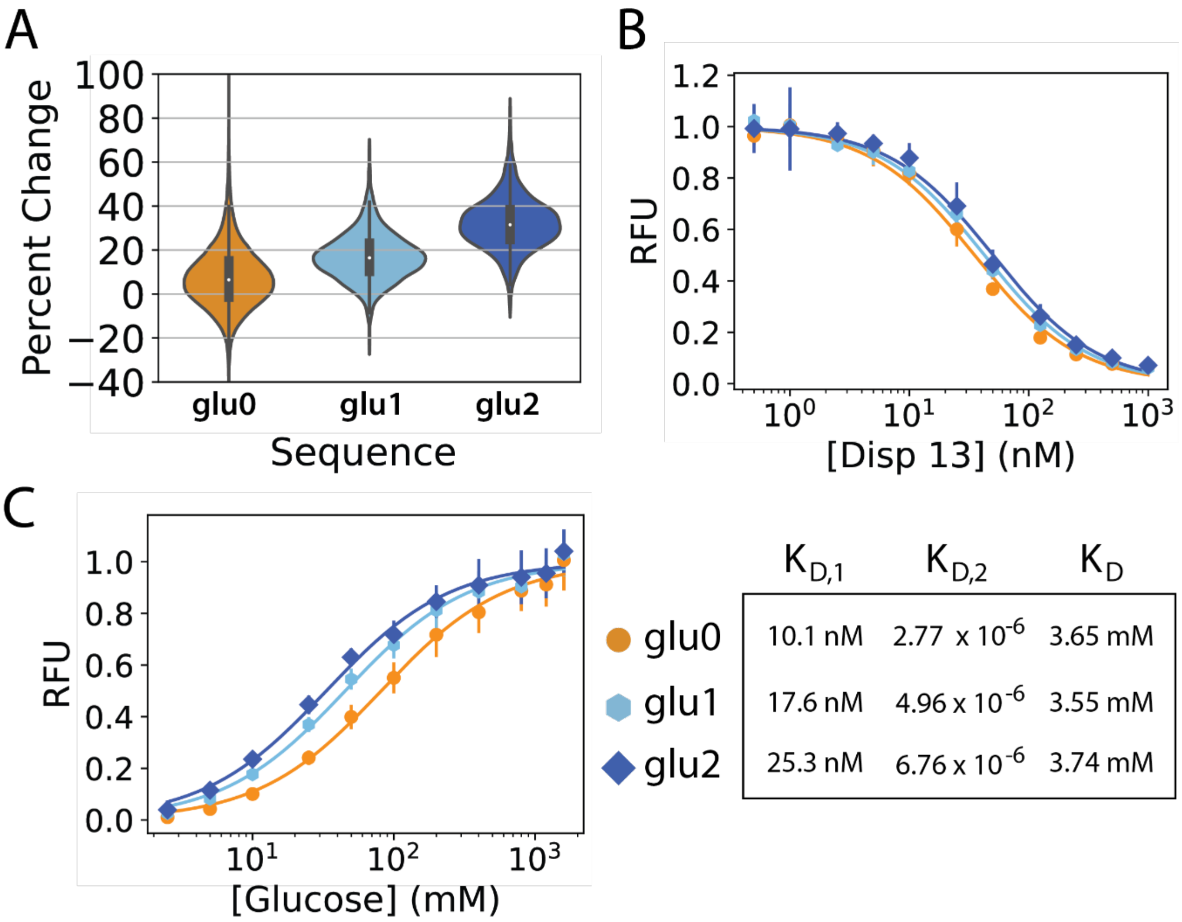


| | $K_{D,1}$ | $K_{D,2}$ | $K_D$ |
|---|---|---|---|
| glu0 | 10.1 nM | $2.77 \times 10^{-6}$ | 3.65 mM |
| glu1 | 17.6 nM | $4.96 \times 10^{-6}$ | 3.55 mM |
| glu2 | 25.3 nM | $6.76 \times 10^{-6}$ | 3.74 mM |

**Supplementary Figure 5**: $K_D$ of parent aptamer glu0 and mutants glu1 and glu2 in HEPES buffer. **A**) Violin plots show percent changes in HEPES buffer from N2A2 analysis. The thick, dark bars in the center of the violin plot show the interquartile range, and the white dots marks the median percent change. $K_D$ values were then measured using a plate-reader assay, and calculated as the ratio of **B)** $K_{D,1}$ and **C)** $K_{D,2}$. The points represent the mean and the bars represent the standard deviation of three measurements. Final values for each calculated $K_D$ are shown in the table at lower right.
